# The Motion Sensitivity and Predictive Utility of Different Estimates of Inter-regional Functional Coupling in Resting-state Functional MRI

**DOI:** 10.1101/2025.07.13.664614

**Authors:** Kane Pavlovich, James C. Pang, Alex Fornito

## Abstract

Numerous methods exist for quantifying statistical dependencies, termed functional coupling (FC), between regional brain activity recorded with resting-state functional magnetic resonance imaging (rs-fMRI). However, their efficacy in mitigating the effects of known sources of noise, such as those induced by participant head motion, and in augmenting effect sizes for brain-wide association studies (BWAS), remains unclear. Here we compared 10 different measures of FC, including correlations, partial correlations, coherence, mutual information, and partial information decomposition, and one measure of effective connectivity (EC; regression dynamic causal modelling), across two independent datasets comprising a total of 1,797 participants (867 males). Each method was evaluated for its ability to mitigate motion-related confounds in FC/EC estimates and for its utility in predicting 94 behavioural measures, as assessed via cross-validated kernel ridge regression. Our analyses showed that EC was most resistant to motion artifacts but had the weakest behavioral predictions. Conversely, traditional correlation-based methods showed the highest sensitivity to motion, but offered the strongest behavioral prediction across most domains and datasets. Nonetheless, relative differences in predictive accuracies were small, indicating that the use of different FC or EC metrics in rs-fMRI does not significantly impact BWAS effect sizes.

---

The development of resting-state functional magnetic resonance imaging (rs-fMRI) marked a paradigm shift in neuroscience, revealing that the brain’s spontaneous fluctuations reflect the intrinsic organization of large-scale neural networks (Biswal et al., 1995; Ogawa et al., 1998). Different measures of inter-regional functional coupling (FC), which quantify statistical dependencies in these fluctuations between brain regions (Friston, 1994), have emerged as popular ways of investigating the architecture of such networks. FC, as commonly quantified using inter-regional signal correlations, exhibits systematic individual differences that correlate with variations in cognition and behaviour (Finn et al., 2015), motivating the rise of brain-wide association studies (BWAS) that use whole-brain FC to predict measures of cognitive ability, psychopathology, and personality (Gratton et al., 2018; Kang et al., 2024; Cheng et al., 2015).

Despite the popularity of FC, recent work has shown that its effect sizes in BWAS are small (i.e., mean correlation of *r* = 0.1) and generally only reproducible in samples exceeding *N* = 2000 (Marek et al., 2022). One possible reason for such small effects is that FC in human studies is often quantified with fMRI, which provides an indirect measure of brain activity and is notoriously noisy (Logothetis, 2008; Buxton, 2013; Power et al., 2012), limiting one’s ability to accurately measure behaviourally relevant brain activity. Several studies have attempted to address this issue by using various preprocessing pipelines to increase the signal-to-noise ratio of FC measures, such as Independents Components Analysis (ICA) based denoising, regression of non-neuronal signals, or the removal of contaminated fMRI volumes (Parkes et al., 2017; Ciric et al., 2017; Parkes et al). However, when applied in BWAS, these tools often fail to substantially increase effect sizes (Pavlovich et al., 2024; Li et al., 2019).

One consideration that has received less attention is how FC is quantified after the data have been processed. There are many different methods for quantifying time series dependencies (Cliff et al., 2023). By far, the most popular technique for quantifying FC is the product-moment correlation between regional fMRI time series, but this measure has several limitations, including a sensitivity to indirect, “third-party” effects (i.e., if region *i* is correlated with *j*, and *j* with *k*, then *i* and *k* will generally show some non-zero correlation, even if they are not truly communicating with each other; Zalesky et al., 2012); an insensitivity to non-linear dependencies; an inability to examine dynamics unfolding at distinct frequencies; and an agnosticism with regards to causal influences (i.e., correlations are undirected measures that do not distinguish whether region *i* causally drives activity in region *j* or vice-versa).

Alternative measures address these limitations to varying extents. For instance, partial correlation coefficients can mitigate third-party effects (Marrelec et al., 2006) but can be prone to their own biases, such as poor estimation of FC in dense networks or in cases of multicollinearity (Zalesky et al., 2012). Mutual information (MI)-based metrics can capture linear and non-linear dependencies (Escudero et al., 2009; Mahadevan et al., 2021), and coherence-based measures can distinguish frequency-specific interactions (Bowyer, 2016; Grinsted et al., 2004). More recently, partial information decomposition (PID) has enabled separation of redundant (shared) and synergistic (emergent) components of FC (Luppi et al., 2021; Luppi et al., 2024). In contrast to FC, dynamic causal modelling (DCM) has been used to infer causal interactions between neuronal populations (Stephan et al., 2010; Friston et al., 2003; Friston et al., 2009), termed as effective connectivity (EC). A recent variant of DCM, called regression DCM (rDCM), offers a method for quantifying EC that is scalable to whole-brain networks (Friston et al., 2003; Frässle et al., 2021).

The multitude of approaches for quantifying coupling between fMRI time series raises questions about their comparative utility and validity. Seminal work by Smith et al. (2011) compared 20 different methods with respect to ground-truth simulations and found that covariance-based methods, such as partial correlations and inverse regularised partial correlations, can recover true network interactions with the greatest accuracy. Follow-up simulation studies have argued that granger causality more effectively recovers regional coupling in fMRI data (Wang et al., 2014; Seth et al., 2013), although regional variations in haemodynamic lags complicate inference about causal interactions in the absence of an appropriate model of neurovascular coupling (Friston et al., 2009). Subsequent studies have extended this approach, comparing the efficacy of different metrics in reducing motion effects on FC estimates in non-simulated rs-fMRI data (Mahadevan et al., 2021; Mohanty et al., 2020). These studies have found that partial correlation-based methods show the lowest levels of residual motion in final FC estimates.

Despite these advances, it remains unclear whether certain FC measures are more sensitive for extracting behaviourally relevant aspects of neurophysiological coupling that may improve BWAS effect sizes. Liu et al. (2024) examined the efficacy of 239 different FC estimations in BWAS predictions and found that covariance-based correlation and partial correlations provided the best performance. However, their analysis focused exclusively on relative effect sizes between metrics and did not report absolute effect sizes. They also did not consider the degree of motion contamination in each FC estimate.

To address this gap, we evaluated 10 different FC measures and one EC measure in terms of the degree to which they: (a) remained sensitive to the contaminating effect of head motion; and (b) influenced the accuracy of models in predicting various aspects of cognition, personality, and psychopathology. We performed the analysis in two independent samples, the Human Connectome Project (HCP) and the Adolescent Brain Cognitive Development study (ABCD) allowing us to determine whether any specific approach offered a reproducible and generalizable way of mitigating noise while maximising BWAS effect sizes.

## Methods

### 1. Participants and data

We used resting-state fMRI data collected as part of the Human Connectome Project (HCP; *N*=1,200) (Van Essen et al., 2013) and the Adolescent Brain Cognitive Development study (ABCD, *N*=11,875) (Karcher & Barch, 2001).

The HCP dataset was acquired on a customized Siemens 3T Skyra at Washington University using a multiband sequence. Structural data were acquired with a T1-weighted (T1w) MPRAGE (TR = 2400 ms, TE = 2.14 ms, flip angle = 8°). Resting-state fMRI data were acquired using a gradient-echo echo planar imaging (EPI) (TR = 720 ms, TE = 33.1 ms, flip angle = 52°) across 1200 frames. Behavioural data were primarily collected using the National Institutes of Health (NIH) toolbox that assessed neurobiological and behavioural function (Gershon et al., 2013), as well as supplementary tests that covered fluid intelligence (Penn Matrix Analysis test, Gur et al., 2010) and personality (NEO personality inventory, Costa & McCrae, 2000). Details on the specific behavioural items used in this study can be found in Supplementary Table 1. Further details on the full list of HCP behavioural measures can be found in Van Essen et al. (2012).

The ABCD dataset includes scans acquired from 21 different sites. To minimize the influence of site effects (Nielson et al., 2018; Focke et al., 2011) and for computational tractability, we used baseline data from the two largest sites (N=1,090, see section 2.3; with data harmonised with ComBat, see section 3), in which scans were acquired on two separate 3T Siemens Prisma Fit scanners using a multiband sequence. Structural data were acquired with a T1w MPRAGE (TR = 2500 ms, TE = 2.88 ms, flip angle = 8°). Resting-state fMRI data were acquired using multiband EPI sequences (TR = 800 ms, TE = 30 ms, flip angle = 52°) for a minimum of 375 frames (Casey et al., 2018; Hagler et al., 2019). Behavioural data were collected using a range of tools, including the NIH toolbox to assess cognitive functioning (Gershon et al., 2013) and the Child Behaviour Checklist (CBCL) to assess behavioural and emotional problems (Achenbach & Rescorla, 2001). Details on the specific behaviours used in this study can be found in Supplementary Table 2. The full list of measures employed by the ABCD can be found in their data dictionary (https://data-dict.abcdstudy.org).

### 2. Image Preprocessing

#### 2.1 Structural Image Preprocessing

In both HCP and ABCD datasets, skull-stripped T1w images in MNI152 space were segmented into white matter (WM), cerebrospinal fluid (CSF), and grey matter (GM) probability maps using the new segment routine from the Statistical Parameter Mapping software v8.0 (SPM8). WM and CSF probability maps were then binarized to create tissue-specific masks. As suggested by Power et al. (2017), WM masks were eroded five times and CSF masks twice to avoid any overlap with the GM signal. Following erosion, if any mask had less than 5 voxels present the previous erosion cycle was selected as the final mask.

#### 2.2 HCP Functional Image Processing

Previous studies have evaluated different preprocessing pipelines based on their ability to maximize effect sizes in BWAS. Although no single optimal approach has been identified, the available evidence suggests that combining movement regressors, manually trained ICA denoising, and global signal regression (GSR) yields good performance across different datasets (Pavlovich et al., 2024; Li et al., 2019), including in the HCP (Pavlovich et al., 2024). We therefore adopted this preprocessing approach for our study.

The HCP data were initially preprocessed according to the HCP minimal processing pipeline version 3.21 (Glasser et al., 2013). Specifically, the T1w images were corrected for gradient distortion, bias corrected, aligned to participants’ T2w images, and registered to the MNI152 2009 non-linear asymmetric template using an affine transformation, before brain extraction (Glasser et al., 2013).

fMRI volumes underwent a gradient distortion correction before realignment to a single band reference image. Image distortion was corrected using reverse coded spin echo maps and each corrected 3D image was registered to the MNI152 2009 non-linear asymmetric template using a non-linear transformation obtained using single spline interpolation. The data were then intensity normalized relative to the value of 1000. Noise-related components were removed using ICA-based X-noiseifier (ICA-FIX; Griffanti et al., 2014). Then, six head motion parameters—along with their temporal derivatives and quadratics—were regressed from the time series. These steps were applied by the HCP data processing team. Further details of each step can be found in Glasser et al. (2013).

We then used linear regression to remove the shared variance with averaged signals from WM, CSF, and entire brain (i.e., the global signal, GS), along with their temporal derivatives from the masks described in section 2.1. The data were then bandpass-filtered (0.008–0.08 Hz) using a Fourier-based rectangular filter. Regression was performed using scripts from the CBIG repository (https://github.com/ThomasYeoLab/CBIG), while filtering was implemented via the *rest_IdealFilter* function (Jia et al., 2019). GSR remains debated in resting-state fMRI preprocessing, but we included it due to evidence that it enhances BWAS effect sizes (Li et al., 2019; Pavlovich et al., 2024).

#### 2.3 ABCD Functional Image Processing

The ABCD data were initially processed using ABCD-specific preprocessing pipelines. First, the T1w images underwent gradient distortion correction via scanner-specific nonlinear transformations, intensity inhomogeneity correction using B1-bias field estimation, and alignment of T1w images to T2w images using mutual information (Hagler et al., 2019). We then aligned T1w images to the MNI152 2009 non-linear asymmetric template using *antsRegistrationsyn.sh* (Avants et al., 2008) and performed brain extraction using *antsBrainExtraction.sh* (Avants et al., 2011).

Functional images were initially processed by the ABCD consortium using their standardized pipeline (Hagler et al., 2019). This included motion correction through volume-wise registration to the first image, B_0_ distortion correction via a reverse polarity approach, gradient nonlinearity distortion correction, and co-registration to structural T1w images using mutual information.

Following this preprocessing, we removed the first 9 volumes from each run to account for magnetic field stabilization. Subsequently, functional images were aligned to the MNI152 2009 nonlinear asymmetric template using transformation matrices derived from T1w image registration (*antsApplyTransforms*; Avants et al., 2011), with brain extraction performed using masks generated during the T1w image processing. For denoising, we trained ICA-FIX on a subset of 24 subjects (balanced by sex and site), achieving a classifier performance with a 85.8% true positive and 80.3% true negative rates (Griffanti et al., 2014).

This classifier was then applied to remove noise components from all remaining data. Additional nuisance regression was performed using scripts from the CBIG repository, incorporating: (1) 24 motion parameters (6 rigid-body parameters, their derivatives, and squared terms); and (2) WM, CSF, and GS signals with their derivatives. Finally, the data were bandpass filtered (0.008–0.08 Hz) using *rest_IdealFilter* (Jia et al., 2019).

### 3. Estimation of Functional Coupling and Effective Connectivity

Prior to FC estimation, all functional images were parcellated by calculating the mean time series for each of 300 cortical regions defined using a widely used parcellation (Schaefer et al., 2018) that shows strong performance in various benchmarking tests (Schaefer et al., 2018; Yan et al., 2023; Pang et al., 2025). We then calculated 11 different FC/EC metrics for evaluation (Table 1, Figure 2A). Following FC/EC estimation, ComBat was used for site harmonization in the ABCD dataset (Johnson et al., 2007; Fortin et al., 2018). ComBat was applied to all edges in each connectivity estimator, using parametric corrections, with age and sex included as covariates to preserve biologically relevant differences in connectivity estimates (Fortin et al., 2018).

**Table 1.**
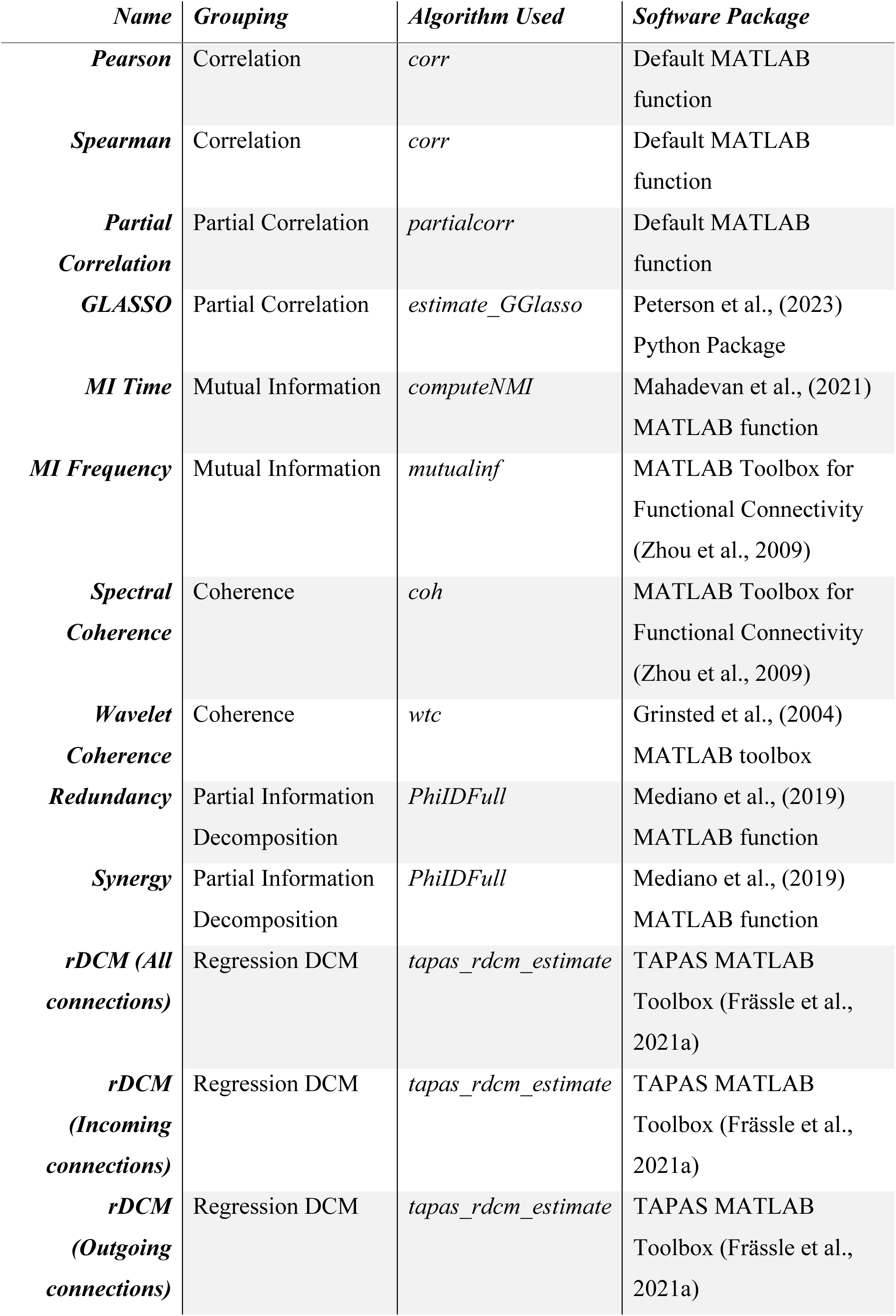
Summary of functional connectivity measures and implementation details.

#### 3.1. Correlations

The classical product-moment correlation is the most popular measure of FC. It quantifies linear temporal dependencies between regional time series, providing a straightforward interpretation of physiological coupling between regions (Smith et al., 2009; Van Dijk et al., 2010). Pearson’s correlation can be defined as:

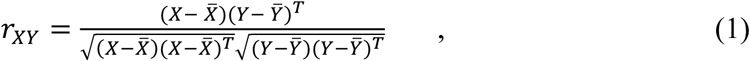

where *X̅* and *Y̅* are the mean values of the signals in regions *X* and *Y*, respectively, and *T* is the transpose of the signal vectors. Spearman’s correlation can be defined as:

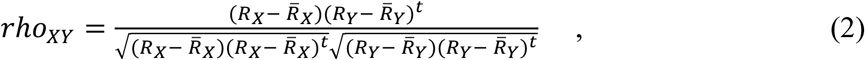

where *R_X_* and *R_γ_* are the rank-transformed time series of *X* and *Y*, respectively (with tied ranks resolved by averaging), and *R̅_X_* and *R̅_γ_* are the mean ranks of *R_X_* and *R_γ_*. Pearson’s and Spearman’s correlation matrices were estimated using the *corr* function in MATLAB. Fishers *r*-to-*z* transformation was applied after estimation to normalise the correlation distribution.

#### 3.2. Partial Correlations

We computed partial correlation matrices for each participant’s parcellated time series to estimate direct FC estimates while controlling for third-party effects. Partial correlations quantify the unique relationship between each pair of regions by regressing out the influence of all other regions in the network (Marrelec et al., 2006; Smith et al., 2011), defined as:

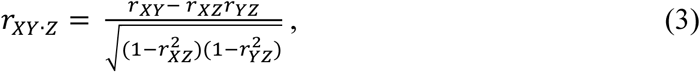

where *Z* refers to a set of variables to be controlled for (i.e., all other regions aside from *X* and *Y*), and *r_XZ_* and *r_YZ_* refer to the correlation between regions *X* and *Y* and all other regions, respectively. All analyses were performed using MATLAB’s *partialcorr* function.

We additionally estimated FC using regularized partial correlations, as per the L1-penalized graphical lasso (GLASSO) (Friedman et al., 2008; Peterson et al., 2023). This method computes partial correlations by inverting the covariance matrix of all nodes to obtain the precision matrix (Θ). A sparsity constraint is then imposed on the precision matrix, driving weaker connections to zero while preserving stronger, more meaningful connections, by adding an L_1_ penalty term to the cost function. The sparsity term is given by

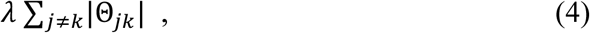

where *j* and *k* are the rows and columns, respectively, of the precision matrix Θ, and *λ* is the hyperparameter controlling the amount of regularisation. The regularised partial correlation coefficients can then be calculated as

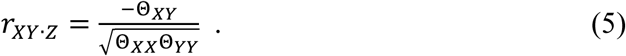

We implemented GLASSO using Python’s *estimate_GGlasso* function. As the optimal regularization strength (*λ*) varies across datasets, we first identified appropriate *λ* ranges for each dataset by running regularization across a wide range of *λ* values in a subset of 20 randomly selected participants. From these initial results, we then narrowed the *λ* range to focus on a tighter range of potentially optimal values centered around those selected in the subset. This ensured that each dataset’s precision matrix was regularised using a λ range specifically tuned to its individual characteristics (Peterson et al., 2023)

#### 3.4. Coherence

We assessed further aspects of frequency-based FC using coherence metrics. Coherence quantifies the synchrony of oscillatory activity between pairs of brain regions at specific frequencies (Gonzalez et al., 2010) and is defined as

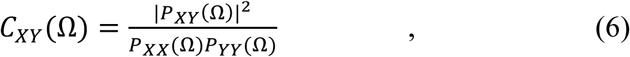

where *P_XY_* (*Ω*) refers to the cross-spectral density between regions *X* and *Y* at frequency *Ω*, and *P_XX_* and *P_YY_* refer to the auto-spectral densities of *X* and *Y*, respectively, at frequency *Ω*.

Spectral coherence was computed using the *coh* function from the MATLAB functional connectivity toolbox (Zhou et al., 2009), which implements Welch’s averaged periodogram method (Welch, 1967). This approach divides the time series into overlapping windows, computes the power spectrum for each window via Fourier transform, and averages these spectra to obtain robust estimates of frequency-specific oscillations. Coherence values were averaged across the 0.008–0.08 Hz band to estimate FC. This averaging approach was used to match prior work utilising averaged frequency estimates (Mahadevan et al., 2021; Mohanty et al., 2020), but we note that multiple distinct frequency bands could be explored using this approach.

To complement this analysis, we assessed time-frequency-resolved functional connectivity using a Morlet wavelet transform (Antonini et al., 1992), implemented via the *wtc* function from the Grinsted MATLAB toolbox (Grinsted et al., 2004). Here, we averaged our connectivity estimates across both time and frequency domains to obtain a time-frequency-resolved estimate of FC, where coherence was averaged across the 0.008–0.08 Hz frequency range.

#### 3.3. Mutual Information

Mutual information (MI) provides a model-free measure of linear and non-linear statistical dependencies between two signals by quantifying their shared information content. In the time domain, the information content of a single region is given its Shannon entropy,

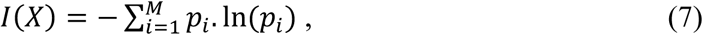

where the regional time series *X(t)* is portioned into *M* bins, and *p_i_* represents the probability of the time series belonging to the *i*-th bin. We computed MI by comparing the Shannon entropy of pairs of time series (Mahadevan et al., 2021), where their joint information is defined as

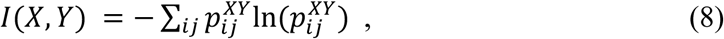

with *I*(*X*, *Y*) being joint distribution between time series *X*(*t*) and *Y*(*t*), and 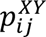 being the joint probability of *X* = *X_i_* and *Y* = *Y_i_*.

Probability distributions for entropy estimation were constructed by binning the time series, with optimal bin numbers determined using the Freedman-Diaconis rule (Freedman & Diaconis, 1981) based on each time series’ interquartile range and sample size. These calculations were implemented using the *computeNMI* MATLAB function (Mahadevan et al., 2021). The MI between pairs of time series was quantified as

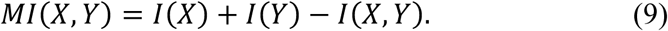

In the frequency domain, MI was derived from partial coherence spectra using the Morlet wavelet transform (Antonini et al., 1992; Grinsted et al., 2004). Specifically, the *mutualinf* function from the MATLAB Toolbox for Functional Connectivity (Zhou et al., 2009) was employed to compute frequency-resolved coupling in the 0.008–0.08 Hz band. This method calculates pairwise coherence from cross-power spectral density estimates (δ_!γ_) and averages coherence values across the specified frequency range (Φ_1_, Φ_2_),

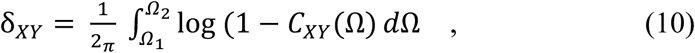

and converts the result to a normalized MI metric (<) using the transformation

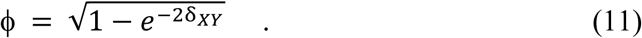

#### 3.5. Partial Information Decomposition

Recently, information decomposition methods have been applied to neural time series data to capture both redundant and synergistic components of network interactions. Here, we applied partial information decomposition (PID) (Mediano et al., 2019; Williams & Beer, 2010) to make this distinction. Redundancy occurs when information is shared across multiple different pairs of regions, and is defined as

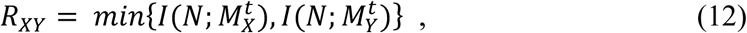

where 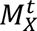 and 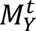 refer to the past states of regions *X* and *Y* at time *t*, respectively, *N* refers to the joint future state of both regions at time *t*+1, and 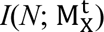 refers to the mutual information between region *X*’s past and its joint future *N*.

Synergy occurs when inter-regional interactions generate emergent properties that are not reducible to individual connections (Mediano et al., 2019; Luppi et al., 2021). Formally,

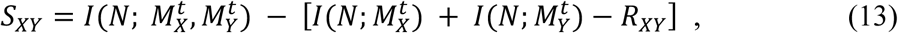

where 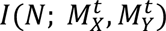 refers to the mutual information between the combined pasts and the future state *N.* PID was implemented using the *PhiIDFull* MATAB function (Mediano et al., 2019), which computes information-theoretic measures across all possible combinations of regional time series. The resulting matrices separately capture redundant signals and higher-order synergistic integration.

#### 3.6. Regression Dynamic Causal Modelling (rDCM)

FC estimates derived from fMRI are limited in many ways, including a sensitivity to measurement noise and a general reliance on undirected measures of coupling, which cannot resolve causal interactions or directions of information flow in the network (Friston, 2011; Friston, 2009; Fornito et al., 2016). Dynamic causal modelling (DCM) is a framework that aims to overcome these limitations by using a Bayesian framework to model the hidden casual neuronal interactions that drive observed fluctuations in haemodynamic signals (Friston et al., 2003). The computational demands of classical DCMs have restricted applications to small networks (typically ≤30 regions) (Novelli et al., 2024), but the recent development of rDCM addresses this limitation by reformulating traditional linear DCM in the frequency domain and recasting it as a special case of Bayesian linear regression. This approach enables significantly faster model inversion, making it possible to estimate effective connectivity in large-scale networks (>100 regions).

Specifically, rDCM approximates causal interactions through Bayesian linear regression in the frequency domain (Frässle et al., 2017) using the likelihood function

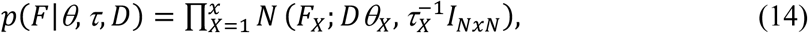

where *F_X_* is the Fourier transform of the temporal derivative of the blood oxygen level dependant (BOLD) signal in region *X*, *D* refers to the design matrix (a representation of endogenous connectivity), *Σ* refers to the afferent connections between regions, *τ_X_* refers to the noise precision parameter for region *X*, and *I_NxN_* is the identity matrix (where *N* denotes the number of data points).

All analyses were implemented using the TAPAS rDCM toolbox in MATLAB (Frässle et al., 2021a; Frässle et al., 2021b). For the present analysis, we examined the full rDCM network alongside EC estimates restricted to either incoming or outgoing connections. This separation allows us to compare the predictive utility of EC and FC predictive models with the same number of features, since FC estimates are symmetric.

### 4. Quality Control Measures

We first evaluated each FC/EC measure in terms of their sensitivity to residual head motion left in the data after pre-processing, which is a major confound in fMRI studies (Power et al., 2012; Van Dijk et al., 2012). To this end, each metric was evaluated using two quality control (QC) measures used extensively to benchmark different preprocessing pipelines (Ciric et al., 2017; Parkes et al., 2017; Pavlovich et al., 2024). The degree to which different FC/EC measures may influence these QC measures has not, to our knowledge, been investigated before.

The two benchmarks we used were Quality-Control FC (QC-FC) correlations and QC-FC distance dependence. QC-FC correlations quantify head motion-related artifacts by computing, for each edge in the FC/EC matrix (44,850 edges for FC; 89,700 edges for EC, across 300 regions), the cross-participant correlation between mean framewise displacement (FD) and FC strength. FD is a measure of the frame-to-frame changes in head motion during a participant’s scan and was derived from each person’s six rigid-body motion parameters (translations and rotations). Following Fair et al. (2020), we first bandpass-filtered these parameters (0.31–0.43 Hz) to remove respiratory artifacts before computing FD as the sum of absolute frame-to-frame differences (Power et al., 2014). The QC-FC matrix thus yields one correlation coefficient per edge, with higher absolute values indicating greater head motion-related information in that connection.

QC-FC distance dependence evaluates how motion artifacts vary with inter-regional distance, as shorter connections are particularly vulnerable to spurious motion-induced coupling (Van Dijk et al., 2012). This dependence was estimated as the Spearman correlation between each edge’s QC-FC value and the Euclidean distance between its regional centroids (Power et al., 2015).

### 5. Participant Exclusions

Following the recommendations from Satterwhite et al. (2013), participants with high levels of in-scanner motion in the HCP dataset were excluded if any of the following stringent motion threshold criteria were met: (i) mean FD > 0.3 mm; (ii) more than 20% of FDs were above 0.2 mm; and (iii) any FD > 5 mm (see also Parkes et al., 2017). Additionally, participants who had any missing behavioural measures were removed from further analysis, such that only participants who had all such measures were included (Figure 1).

**Figure 1.**
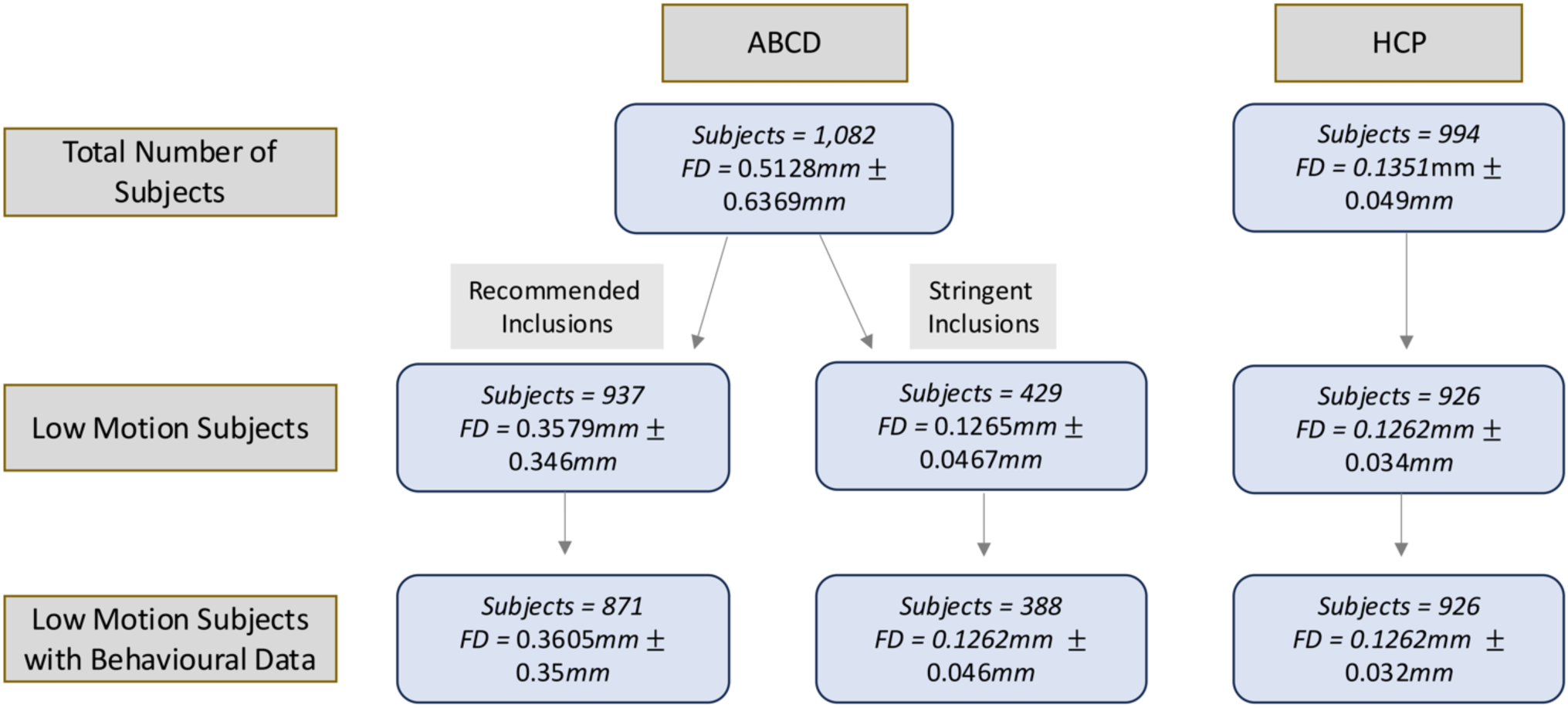
Participant exclusions based on FD and censoring exclusions across the ABCD and HCP datasets. The mean and standard deviation of the FDs across participants are shown.

Given the elevated motion levels characteristic of developmental populations, we performed parallel analyses in the ABCD dataset using two distinct inclusion criteria: (i) the stringent motion threshold based on FD as applied to the HCP data; and (ii) a more lenient quality-controlled subset comprising all participants who passed the standard ABCD fMRI quality assurance protocols (Hagler et al., 2019).

Following these exclusions, our final sample comprised 926 individuals (423 males) in the HCP data, and 388 individuals (177 males) and 871 individuals (444 males) in the ABCD data after stringent and standard exclusion, respectively.

### 6. Behavioural Prediction

#### 6.1. Behavioural Measures

The HCP and ABCD datasets collectively provide comprehensive behavioral phenotyping, with measures spanning cognitive and personality domains in both cohorts, and psychopathology related variables in the ABCD. For our analyses, we selected 58 behavioural measures from HCP (encompassing cognition and personality; Supplementary Table 1) and 36 from ABCD (encompassing cognition, and psychopathology; Supplementary Table 2). Behaviours were selected so that they spanned a large range of both cognitive and personality/psychopathology-related measures across datasets.

Prior to prediction, we regressed age, sex, and FD from each behavioural measure (Siegel et al., 2017; McCarthy et al., 2012; Lucas & Donnellan, 2009). The regression coefficients were calculated from the training set and then applied to the test set to avoid leakage between train and test sets in the prediction model (see below) (Chyzhyk et al., 2022).

#### 6.2. Prediction procedure

To evaluate how different FC/EC estimation methods influence BWAS effect size estimates, we employed a 20-fold cross-validated Kernel Ridge Regression (KRR) (Li et al., 2019; Kong et al., 2019). KRR was chosen as it offers superior or comparable prediction performance relative to other more complex methods such as deep neural networks (He et al., 2020). Separate models were used for FC/EC matrix and each of the 58 behavioural scores in the HCP dataset and 36 behavioural scores in the ABCD dataset. Members from the same family were included in the same test/train fold to avoid train-test predictions across families.

KRR uses an L2 regularisation parameter, which adds a penalty that is equivalent to the sum of the squared values of the weights to the loss function. By doing so, it controls the trade-off between low training error and low validation error, which mitigates overfitting. Following past work (Li et al., 2019, He et al., 2020, Kong et al., 2019; Pavlovich et al., 2024), we selected the value of this regularisation parameter using a repeated nested cross-validation procedure with an outer 20-fold cross-validation and an additional 20-fold inner cross-validation within each outer fold to identify the optimal value for the regularization parameter. This process was repeated 20 times to ensure robust estimation. Prediction accuracy, defined as the correlation between the true and predicted scores, was used as our performance metric (Kong et al., 2019).

Both rDCM and GLASSO impose sparsity constraints on the resulting EC/FC matrices. This constraint may bias the predictive models, since those relying on rDCM or GLASSO estimates will incorporate fewer features than models using FC estimates with no sparsity constraint. We therefore conducted supplementary analyses in which all other FC matrices were thresholded to match the sparsity of the GLASSO-derived matrices (which imposes a higher sparsity constraint compared to rDCM). Specifically, for each participant, we retained only the top *k* connections by weight, where *k* corresponds to the number of connections preserved in the GLASSO output.

### 7. Data and Code Availability

Neuroimaging and behavioural data from both datasets are available at the following links: HCP (https://www.humanconnectome.org/study/hcp-young-adult/document/1200-subjects-data-release) and ABCD (https://nda.nih.gov/study.html?id=2313). For this study we used the baseline data from the 5.1 release of the ABCD dataset.

All code used in our analysis, including links and references to codes used by others, can be found at https://github.com/kanepav0002/Coupling-Estimation-Prediction

## Results

### 1. Subject Exclusions

In the full samples, mean FD was higher in the HCP dataset than in the ABCD dataset. After applying stringent motion exclusion criteria in the ABCD dataset, mean FD values were comparable across retained participants in both datasets (HCP: *M* = 0.1262 ± 0.034 mm; ABCD: *M* = 0.1265 ± 0.0467 mm).

### 2. Similarities between measures

We considered 11 distinct connectivity measures (Figure 2A), each capturing unique aspects of neuronal coupling in rs-fMRI. Figure 2B shows the pattern of inter-correlations between these measures. While some of these measures share overlapping information, others provide complementary insights. Specifically, in the HCP dataset, mutual information, coherence, and redundancy-related measures exhibit strong intercorrelations (Pearson’s *r* > 0.85; Figure 2B), indicating shared variance. In contrast, synergy and rDCM-derived connectivity matrices demonstrate only moderate to weak correlations with all other measures (Pearson’s *r* < 0.65; Figure 2B), suggesting they capture novel aspects of interregional coupling. Results were similar in the ABCD dataset (in both the stringent and recommended subset) where mutual information, coherence, and redundancy-related measures also exhibited strong intercorrelations (Pearson’s *r* > 0.83; Figure 2B), and synergy/rDCM derived matrices demonstrated weak to moderate correlations with all other measures (Pearson’s *r* < 0.72).

**Figure 2.**
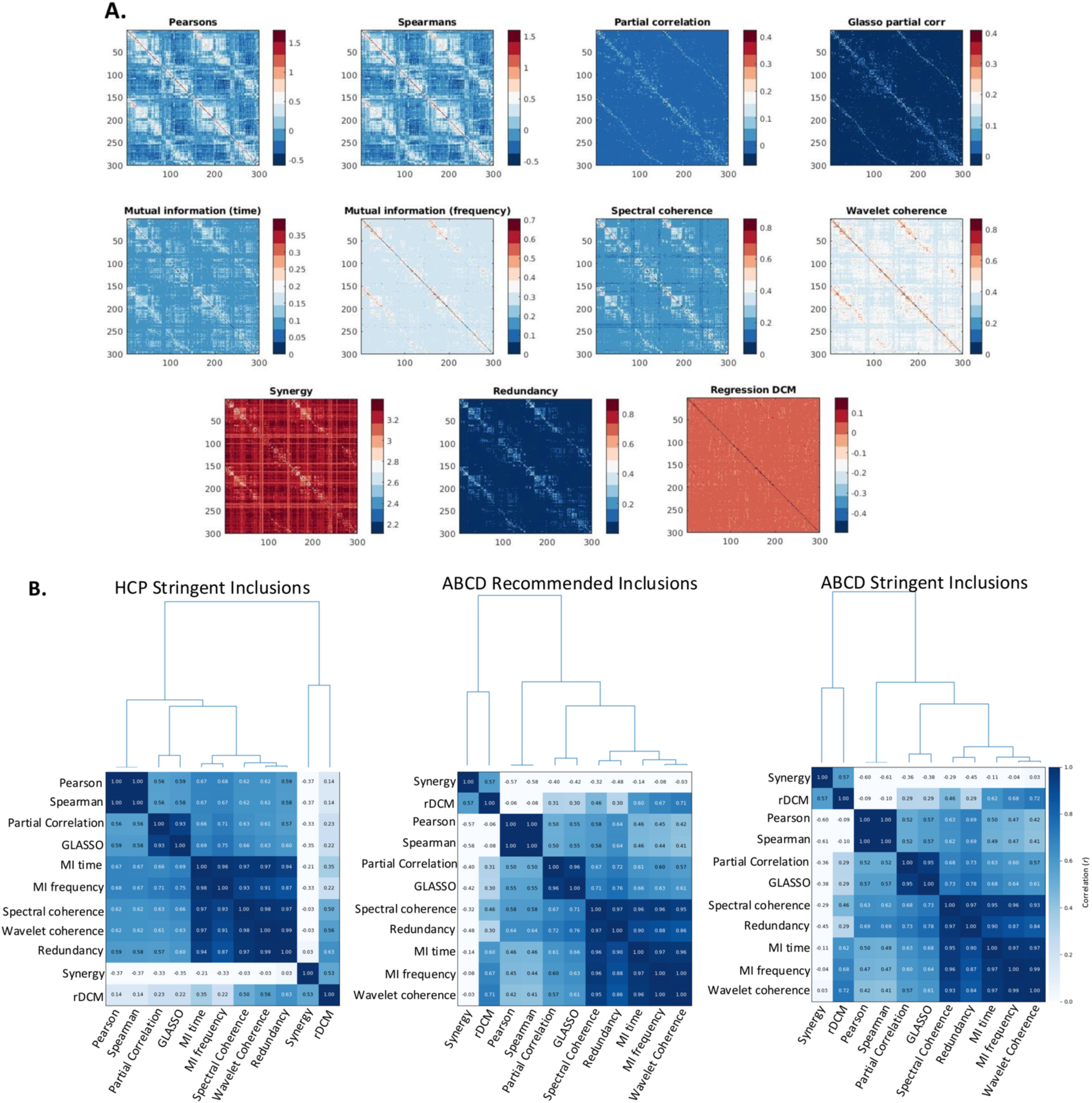
**A.** Group-averaged FC and EC matrices obtained with each of the 11 estimation methods in the HCP dataset after stringent motion exclusions had been applied. **B.** Heatmap of pairwise Pearson correlations among mean FC matrices in the HCP dataset after stringent motion inclusions, and in the ABCD dataset with both stringent and recommended subject inclusions. Metrics are arranged by similarity according to hierarchical clustering using average linkage.

### 3. QC-FC Correlations

We first evaluated the residual motion artifacts in each FC/EC matrix by examining QC-FC correlation distributions across both datasets, where less residual motion is characterized by distributions centred at zero with minimal dispersion. In the HCP dataset, all metrics were approximately centred on zero and had similar variance (Figure 3A). The variance of the rDCM QC-FC distribution was the smallest and the variance of Pearson correlation was the highest, indicating that these measures performed best and worst, respectively.

**Figure 3.**
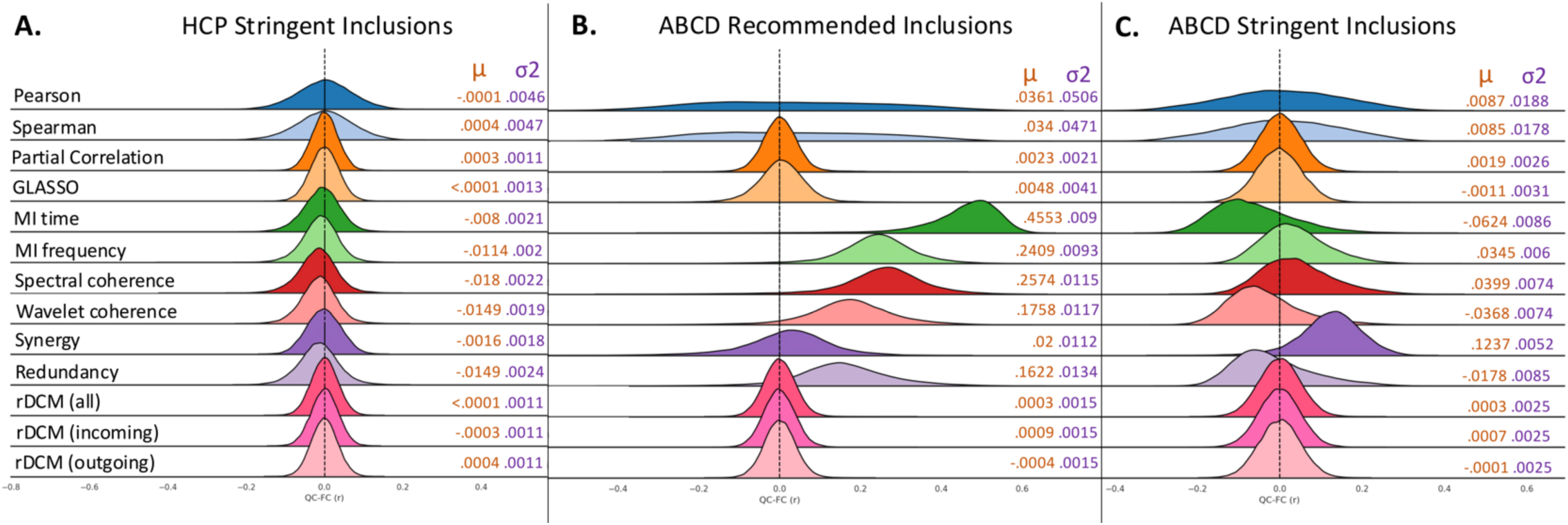
Distribution of QC-FC correlations across all methods for quantifying FC/EC for (**A**) HCP participants meeting stringent motion criteria, (**B**) ABCD participants passing standard quality control inclusion criteria, and (**C**) ABCD participants meeting stringent motion exclusion thresholds. Measures are ordered as: Correlation → Partial correlation → Mutual Information → Coherence → PID → rDCM. Distribution means (μ) are presented in orange text, while population variance (02) is presented in purple text.

In the ABCD cohort defined by recommended inclusions, we observed considerable variability in performance across metrics (Figure 3B). As in the HCP dataset, rDCM showed the strongest performance, with a mean of .0003 and a variance of .0015, which was the lowest of all pipelines. Pearson correlation also performed the worst, showing an almost uniform distribution, indicative of a comparatively large number of high QC-FC values. Spearman correlation showed a similar pattern. MI, coherence, and PID-derived metrics all showed a right-shift in their QC-FC distributions, indicating that these methods are particularly sensitive to motion contamination, with the worst performing being MI in the time domain.

Notably, restricting the ABCD cohort to low motion participants after applying stringent criteria was associated with a drastic mitigation of these effects (Figure 3C), highlighting the importance that judicious inclusion of participants has in mitigating noise within high-motion datasets. EC estimates derived from rDCM once again performed the best, followed closely by partial correlation and GLASSO. Pearson and Spearman correlations were associated with zero-centred distributions with large variance. The right-shift in the QC-FC distributions of MI-frequency, Spectral Coherence, and Redundancy were still present, albeit attenuated in magnitude. For MI time, wavelet coherence, and Synergy, the right-shift observed following less stringent inclusion criteria changed to a left-shift, indicating that these metrics still showed significance motion contamination, albeit with a reversed polarity of association between FC and motion. Controlling the level of sparsity to match the constraints imposed by the GLASSO partial correlation did not change the pattern of overall results observed across all three cohorts (Supplementary Figure 1).

### 4. QC-FC Distance Dependence

We next examined the efficacy of each FC/EC measure in mitigating the distance dependence of QC-FC correlations. In the HCP dataset, all metrics showed similar levels of distance dependence, with correlations ranging between -0.08 < r < 0.05. rDCM was associated with the smallest distance-dependence, whereas MI in the time domain was associated with the largest (Figure 4A). In the ABCD cohort defined by recommended inclusions, distance-dependence showed considerable variation across metrics. The lowest distance dependence was observed for rDCM, closely followed by partial correlation (Figure 4B). Distance dependence was substantially higher for Pearson and Spearman correlations. A similar pattern was observed following the application of stringent inclusion criteria (Figure 4C). Controlling for sparsity did not largely influence the pattern of findings (Supplementary Figure 2).

**Figure 4.**
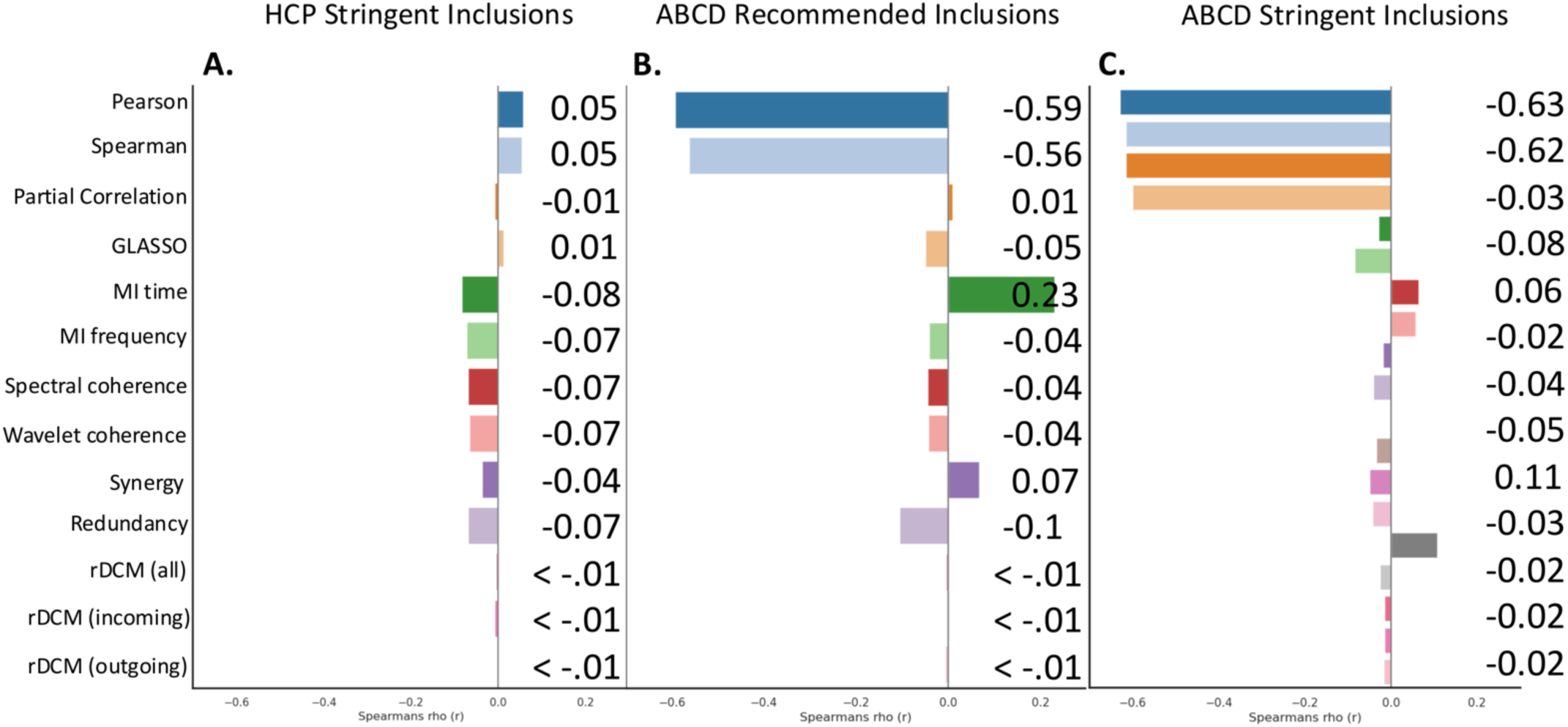
QC-FC distance-dependence correlations across datasets. Bars show the distance dependence value for each connectivity metric for (**A**) HCP participants meeting stringent motion criteria, (**B**) ABCD participants passing standard quality control inclusion criteria, and (**C**) ABCD participants meeting stringent motion exclusion thresholds. Measures are ordered as: Correlation → Partial correlation → Mutual Information → Coherence → PID → rDCM.

### 5. Behavioural Prediction

We evaluated the predictive utility of the FC/EC measures for out-of-scanner behavioural measures (36 in ABCD, 58 in HCP) using separate multivariate KRR models for each metric and behaviour. Prediction accuracy was quantified as the product-moment correlation (*r*) between true and predicted behavioural scores, averaged across cross-validation folds, repetitions, and behaviours (Figure 5). We considered the following three behavioural domains separately: (1) cognitive measures; (2) psychopathology (ABCD only); and (3) personality traits (HCP only; see Supplementary Tables 1–2 for measure-specific inclusions).

**Figure 5.**
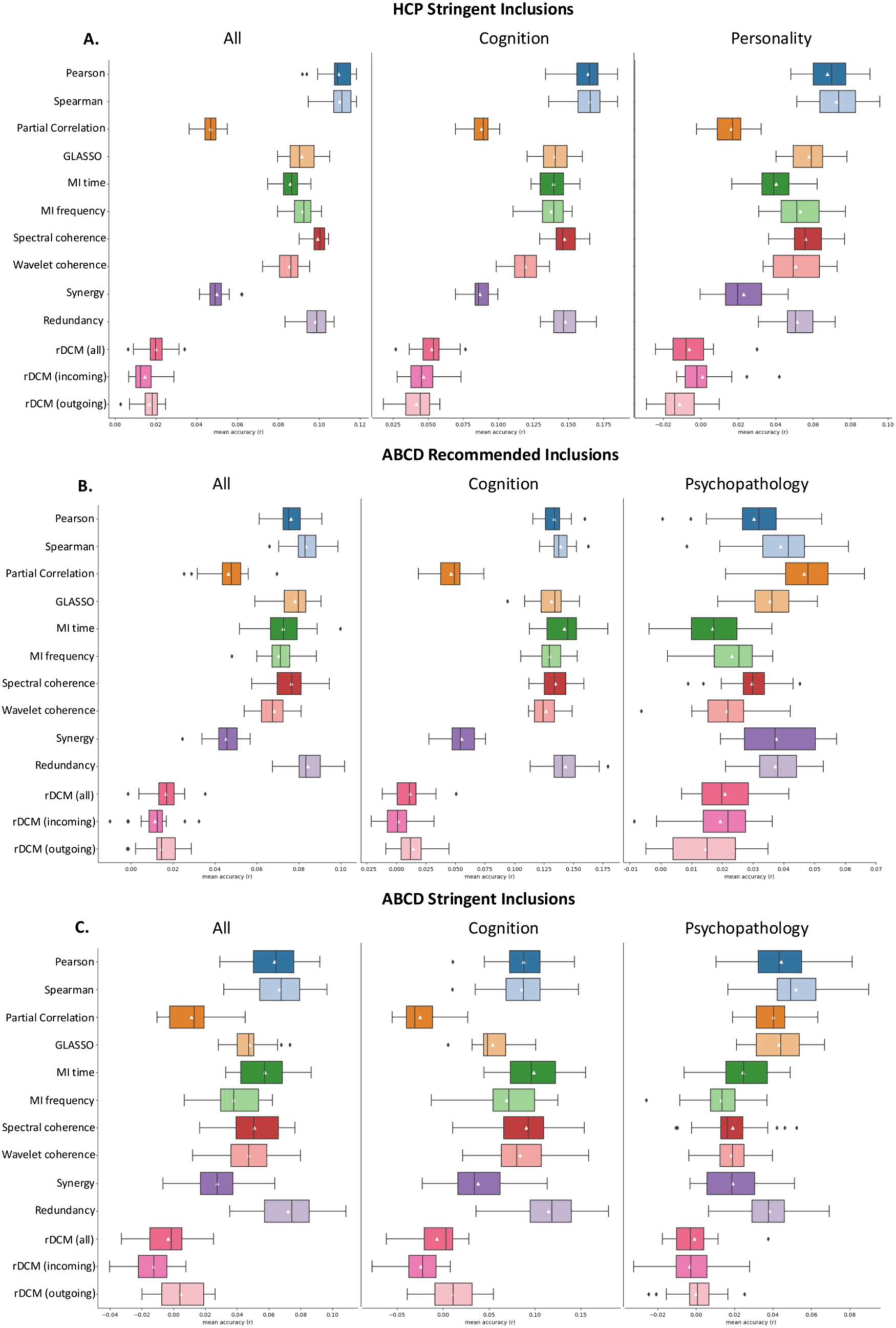
KRR prediction accuracies across datasets. The boxplots show the median and interquartile ranges for accuracies averaged over cross validation folds, repetitions, and behaviours. Mean predictive accuracy for each connectivity measure is shown with a white triangle. Prediction accuracies are shown for (**A**) HCP participants meeting stringent motion criteria, (**B**) ABCD participants passing standard quality control inclusion criteria, and (**C**) ABCD participants meeting stringent motion exclusion thresholds. Measures are ordered as: Correlation → Partial correlation → Mutual Information → Coherence → PID → rDCM

For cognitive measures, the optimal method showed some variation across datasets and participant inclusion criteria. In the HCP dataset (Figure 5A), Pearson and Spearman correlations were associated with the highest predictive accuracies (mean *r* of 0.164 and 0.165, respectively), whereas rDCM was associated with the worst, being essentially unpredictive of behaviour (mean *r* = 0.053). Partial correlation also showed poor performance (mean *r* = 0.088) which was, to some extent, rescued by regularization via GLASSO (mean *r* = 0.14). MI and coherence-based metrics showed similar performance to each other (0.119 < mean *r* < 0.139).

In the ABCD cohort, Redundancy was associated with the best predictive performance, followed closely by Pearson and Spearman correlations. rDCM was again associated with the worst performance. Performance variations were consistent under recommend and stringent exclusion criteria, although effect sizes were slightly higher in the former cohort (e.g., for Redundancy, mean *r* was 0.143 for recommended criteria and 0.115 and stringent criteria) (Figure 5B; Figure 5C).

Similar trends were observed for prediction of personality in the HCP, albeit with smaller effect sizes (i.e., all *r* < 0.1) (Figure 5A). Once again, Pearson and Spearman correlations were associated with the highest accuracies whereas rDCM-based estimates were associated with the lowest accuracy.

For psychopathology in the ABCD sample, Pearson and Spearman correlations yielded the best performance in the stringent sample (Figure 5C). These measures also performed well in the sample defined by recommended inclusions (Figure 5B), although partial correlation performed best in this group. In both cases, rDCM performed worst.

Interpretation of the psychopathology BWAS findings should be tempered by the fact that, in all cases, prediction accuracies for psychopathology were generally low (i.e., all *r*<0.08). The findings were consistent when all matrices were matched to GLASSO for sparsity (Supplementary Figure 3).

## Discussion

This study systematically evaluated 10 FC measures and one EC measure in two independent rs-fMRI datasets across two key dimensions: (1) robustness to motion-related artifact; and (2) predictive power for cognition, personality, and psychopathology. Our results demonstrate that rDCM consistently shows the greatest resilience to motion contamination across all analyses, but is generally associated with lower BWAS effect sizes; conversely, correlation-based estimates showed poor denoising efficacy but often yielded higher BWAS effect sizes. As such, no single connectivity measure universally maximizes denoising efficacy and BWAS prediction performance, with optimal metrics varying across both behavioural domains and datasets.

### Mitigation of motion-related artifacts

Among all FC/EC metrics evaluated, and across both datasets, traditional correlation-based methods performed the worst with respect to minimizing QC-FC correlations and QC-FC distance dependence. This result is consistent with prior evidence that correlations are highly sensitive to outliers and motion artifacts in rs-fMRI data (Power et al., 2012; Mahadevan et al., 2021). We further observed that motion-related contamination of correlation-based FC was more pronounced in the ABCD dataset, which likely reflects the higher levels of in-scanner motion in this cohort (as indicated by FD; Figure 1). Notably, this elevated motion persisted even after matching FD between cohorts, suggesting residual motion artifacts that are not fully captured by the mean FD-based participant exclusion criteria.

Partial correlations dramatically reduced residual motion influences in the ABCD cohort, consistent with prior work demonstrating that they mitigate motion-induced spurious correlations by accounting for shared variance across regions (Liang et al., 2012; Peterson et al., 2023). Evaluations with respect to ground-truth simulations indicate that partial correlations more accurately isolate pairwise interactions between regions than full correlations due to the removal of third-party effects (Smith et al., 2011), although this performance may depend on the size of the underlying network (Zalesky et al., 2012).

While the confounding effects of motion on FC, particularly for correlation-based estimates, have been well-documented (Mahadevan et al., 2021; Mohanty et al., 2020), the impact of motion artifacts on EC estimates such as rDCM remains largely unexplored. Our analysis indicates that rDCM exhibits greater robustness to motion artifacts than other connectivity measures, showing the lowest motion-related variance across both QC-FC correlation and distance-dependence metrics across both datasets. This resilience may arise from rDCM’s focus on inferring direct neuronal interactions through its regression-based framework (Frässle et al., 2017), which more directly models latent neuronal dynamics rather than relying solely on observed BOLD signal correlations.

Information theoretic and coherence-based measures showed a strong sensitivity to the overall levels of motion in the cohort. They performed well in the HCP dataset but showed particularly poor performance in the ABCD cohort defined by recommended inclusion criteria, which was associated with the highest levels of motion. Applying stringent motion exclusion criteria to the ABCD sample attenuated the level of motion-related contamination for these FC estimates, but they were still often associated with non-zero-centred QC-FC distributions.

Together, our findings indicate that all methods perform similarly with respect to motion contamination in high-quality, low-motion data (e.g., the HCP cohort) and that rDCM is highly robust to motion contamination. Pearson and Spearman correlations show significant contamination in high-motion datasets, as do MI and coherence-based estimates of FC. The levels of overall motion in a cohort, and the motion-related exclusion criteria applied prior to analysis, are thus important factors when considering which FC metric may be most appropriate for a given analysis.

### Prediction of behaviour

Any pipeline or measure for quantifying FC/EC must strike a balance between aggressive denoising and preservation of neural signal. Overly aggressive methods may excel in QC-FC metrics simply because they remove both noise and signal. To assess whether behaviourally relevant information is retained, we evaluated how different methods impact predictive accuracy when using FC/EC to predict behavior (Li et al., 2019; Pavlovich et al., 2024).

Our analysis revealed, consistent with this assumed trade-off between denoising efficacy and preservation of neural signals, that FC/EC measures performing well on motion-related denoising are generally associated with lower prediction accuracies. This effect was particularly pronounced for rDCM, wherein all predictive models were essentially uninformative of behaviour. Conversely, models relying on Pearson and Spearman correlations were generally ranked among the most predictive of cognition, personality, and psychopathology in both datasets. This finding suggests that correlation-based FC may better preserve behaviourally relevant information than more sophisticated methods, at the expense of higher levels of residual motion contamination. In fact, it is possible that such preservation is precisely what drives the predictive performance of correlation-based methods, given prior evidence that more aggressive denoising generally reduces correlations between FC estimates and behaviour (Pavlovich et al., 2024). Motion itself is correlated with different behavioural measures, such as somatic and externalising related psychopathology, as well as abstract reasoning and cognitive flexibility (Siegel et al., 2017), which may be driven by an association between individual differences in head motion and trait-level characteristics (e.g., more impulsive people may move more in the scanner). Our findings suggest that successful denoising of motion-related artifacts may also remove this source of behaviourally relevant variance.

In the ABCD dataset, correlation-based methods were outperformed by two approaches: (1) redundancy-based FC (derived from partial information decomposition, PID) for cognitive predictions; and (2) partial correlations for predictions of psychopathology predictions. The latter only occurred in the subset defined by recommended inclusions and not in the stringent inclusion subset, highlighting once again the importance that inclusion criteria have in shaping one’s results. Redundancy-based FC also showed the second highest cognitive prediction accuracies in the HCP dataset. This result suggests that cognitive abilities may show a stronger association with shared pairwise connections between brain regions than with information that reflects emergent network level properties (Luppi et al., 2022). The stronger performance of partial correlations for predicting psychopathology may reflect a stronger association with direct, linear network interactions that persist after controlling for third-party effects.

These interpretations of comparative differences between FC/EC measures must be tempered by the fact that all prediction accuracies were modest, with all *r* < 0.18, and that differences between approaches were also small (e.g., the mean difference across all measures in the HCP dataset was *r* = 0.008). This result implies that there is no “magic bullet” measure that can significantly uncover a measure of FC or EC with sufficient biological validity to dramatically augment BWAS effect sizes. Whether it is possible to identify such a measure remains an open question.

## Limitations and conclusions

We evaluated a representative selection of FC/EC measures spanning time-versus frequency-domain approaches, linear and non-linear coupling estimators, and undirected versus directed/causal interaction models. While our approach covers the most widely used methods in contemporary rs-fMRI research, our analysis was not exhaustive and many other approaches exist for connectivity estimation (Cliff et al., 2023; Liu et al., 2024). Notably, we did not assess approaches that cover dynamic FC (Hutchinson et al., 2013) or traditional DCMs (Friston et al., 2003), which may offer superior trade-offs between motion robustness and behavioral prediction. Future work should systematically compare these advanced methods using the dual evaluation framework (motion resistance versus predictive validity) conducted here. Other approaches for quantifying the preservation of neural signal (e.g., Aquino et al., 2020; Glasser et al., 2019) may be helpful in this regard.

A fundamental challenge in BWAS research lies in the limited reliability and validity of behavioural measures. Widely used summed scale scores (as employed here) provide practical metrics but they often poorly capture their target constructs (Dang et al., 2020; Pavlovich et al., 2025)—a limitation that inherently attenuates brain-behaviour associations (Saccenti et al., 2022). As such, refinement of neural phenotypes alone cannot meaningfully improve BWAS effect sizes without parallel advances in behavioural measurement. Future studies should employ advanced psychometric approaches (e.g., item response theory, classical test theory) to determine whether optimized behavioral phenotyping can increase the upper bounds of detectable brain-behaviour relationships (Tiego et al., 2023). Repeated measures of behaviour in ecological settings may also assist in this endeavour (Nikolaidis et al., 2022).

In summary, our analysis of 10 different FC measures and one EC measure showed that popular correlation-based approaches are associated with the highest sensitivity to motion artifacts while also yielding the most accurate predictions of behaviour. Conversely, rDCM showed strong resilience to contamination by motion but was essentially unpredictive of behaviour. We also found that the relative sensitivity of different measures to motion and behavioural variance depends on the amount of motion present in the data, although variations in the behavioural prediction accuracies of different pipelines were small. These results underscore the need to carefully consider which individuals should be retained for further analysis, and indicate that the optimum method for quantifying FC or EC should be tailored to the particular question at hand.

## Acknowledgements

A.F. was supported by the National Health and Medical Research Council (ID: 1197431) and Australian Research Council (ID: FL220100184).

J.C.P. was supported by the National Health and Medical Research Council (2034000), Monash FMNHS Early Career Postdoctoral Fellowship, and Monash FMNHS Early Career Research Excellence Program.

This research was supported by Monash eResearch capabilities, including M3 (Massive).

