## Supplementary Information for "The Motion Sensitivity and Predictive Utility of Different Estimates of Inter-regional Functional Coupling in Resting-state Functional MRI"

1. *Matched Sparsity analysis.*

Given that both rDCM and GLASSO partial correlations inherently impose sparsity constraints on connectivity matrices, we conducted a supplementary analysis where all other metrics were thresholded to match the sparsity level of the GLASSO-derived matrices. Specifically, for each participant, we retained only the top X% of connections (where X corresponded to the connection density preserved in the GLASSO output). Such that, if GLASSO retained the top 10% of connections in the matrix, we kept the top 10% of the highest connections in the correlation, mutual information (MI), and coherence measures. As PID retains synergistic and redundant components only, we did not control for sparsity in these matrices.

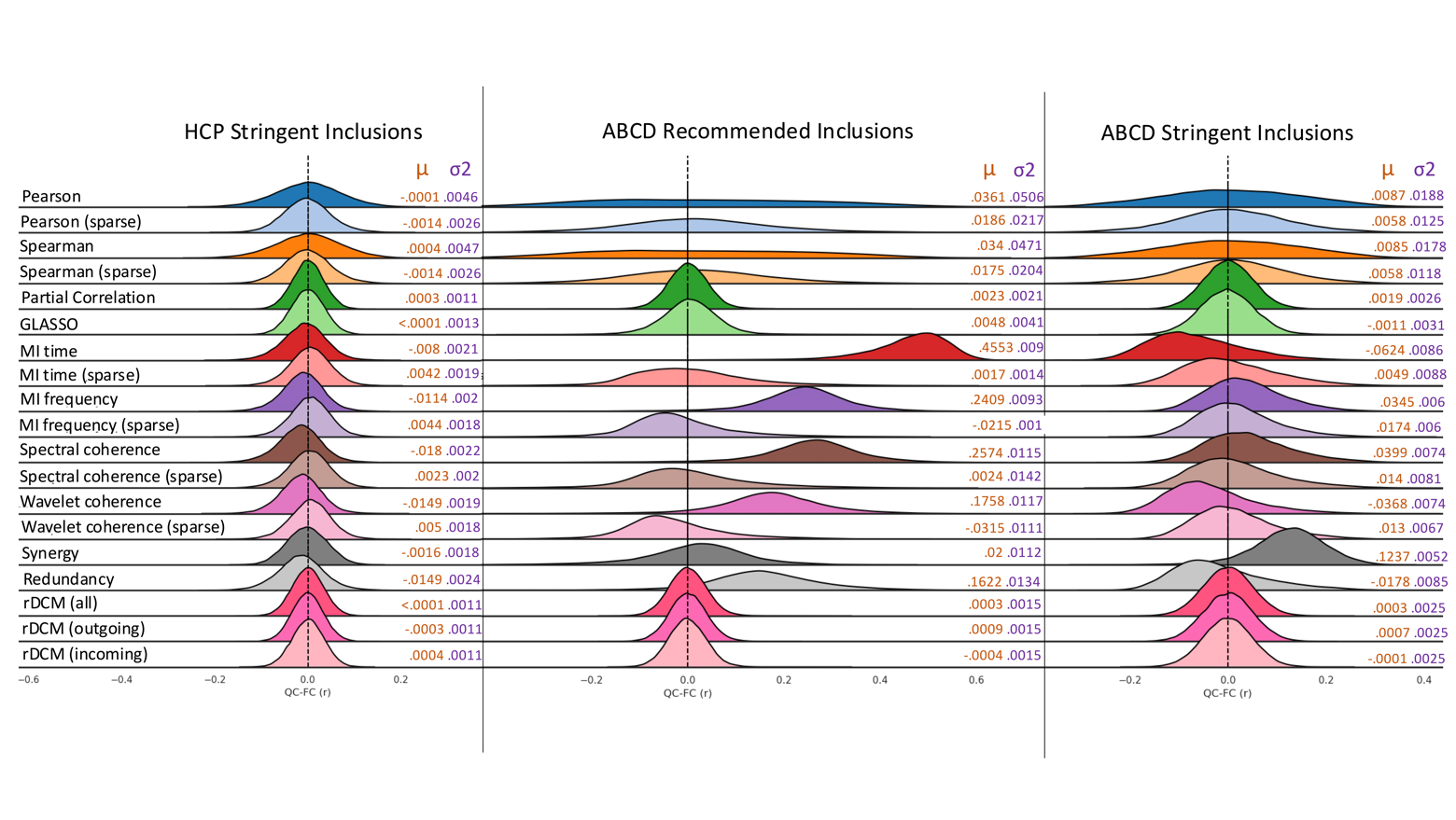

*Supplementary Figure 1.* Distribution of QC-FC correlations across all functional connections presented in the main text and after sparsity constraints were imposed. Distribution plots compare motion profiles for HCP participants meeting stringent motion criteria, ABCD participants passing standard quality control inclusion criteria, and ABCD participants meeting stringent motion exclusion thresholds. Measures are ordered as: Correlation → Partial correlation → Mutual Information → Coherence → PID → rDCM.

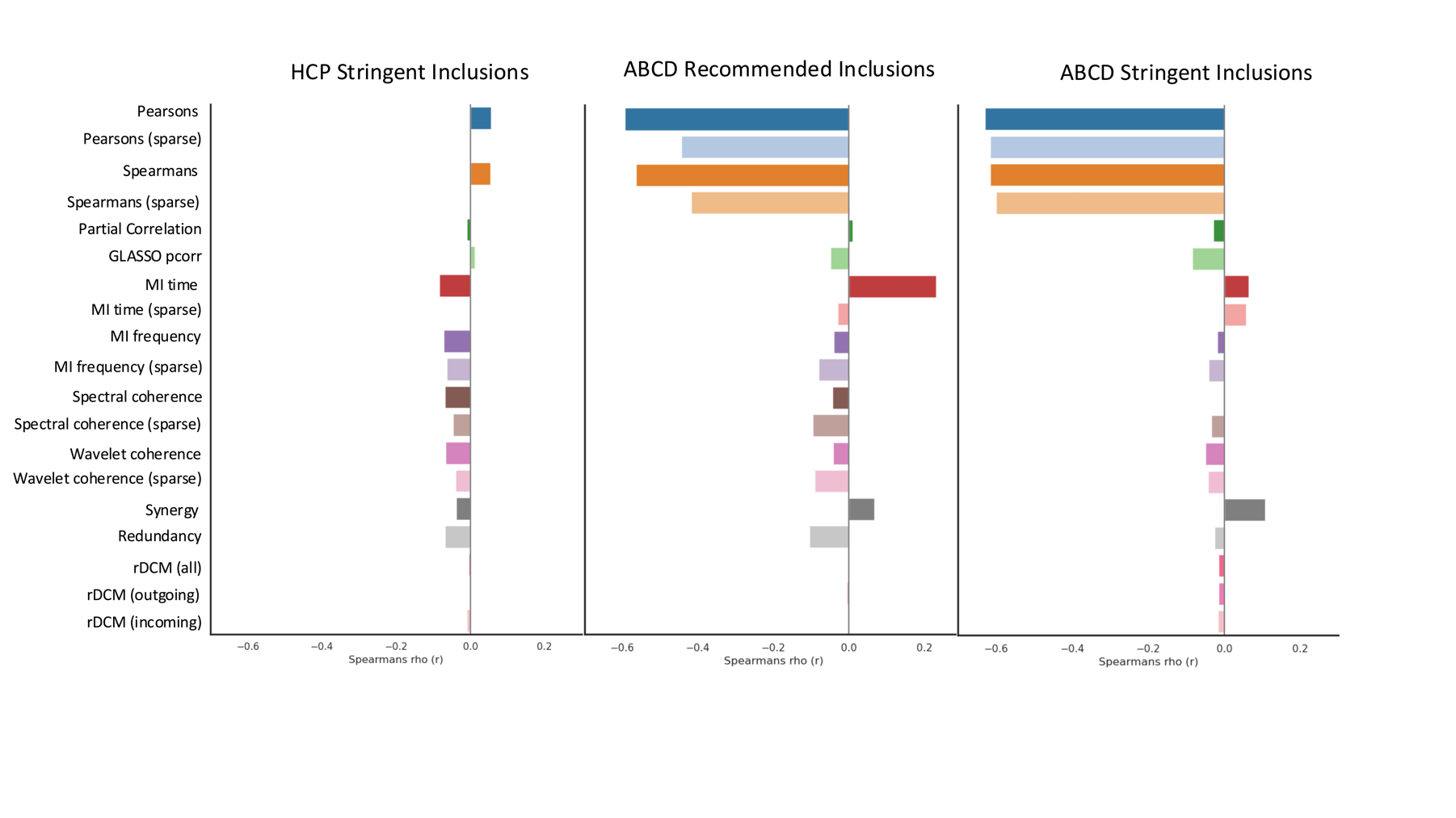

*Supplementary Figure 2.* QC-FC Distance Dependent correlations presented in the main text, including results after sparsity constraints were imposed. Bars show the distance dependent value for each connectivity metric for HCP participants meeting stringent motion criteria, ABCD participants passing standard quality control inclusion criteria, and ABCD participants meeting stringent motion exclusion thresholds. Measures are ordered as: Correlation → Partial correlation → Mutual Information → Coherence → PID → rDCM.

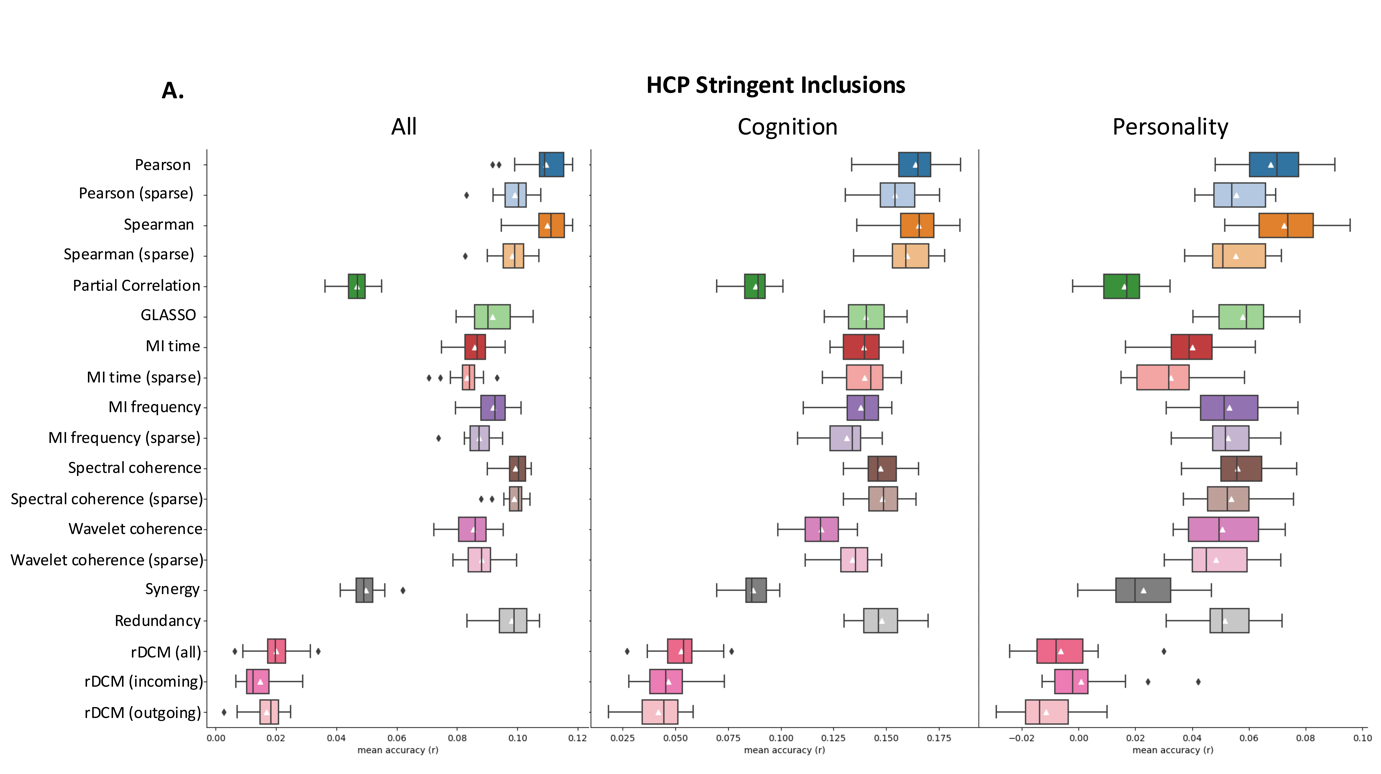

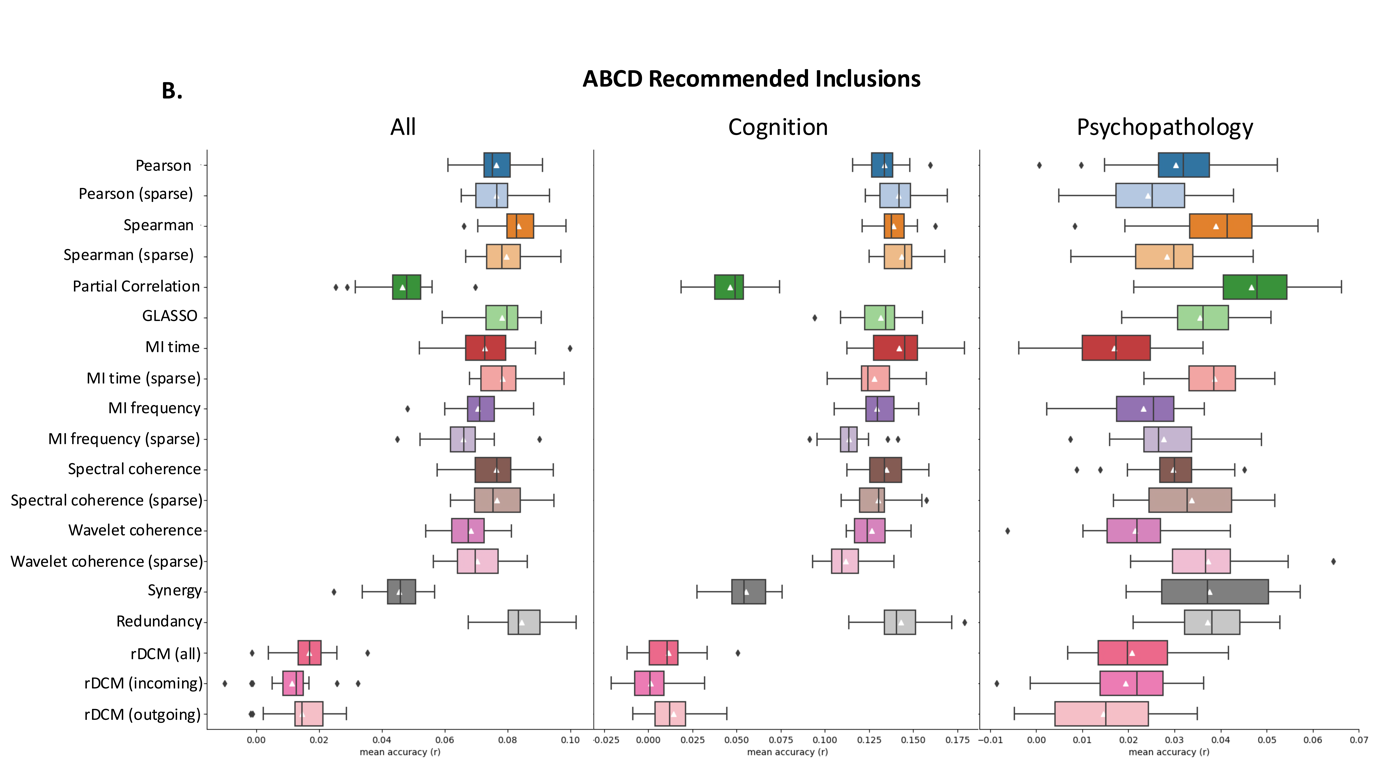

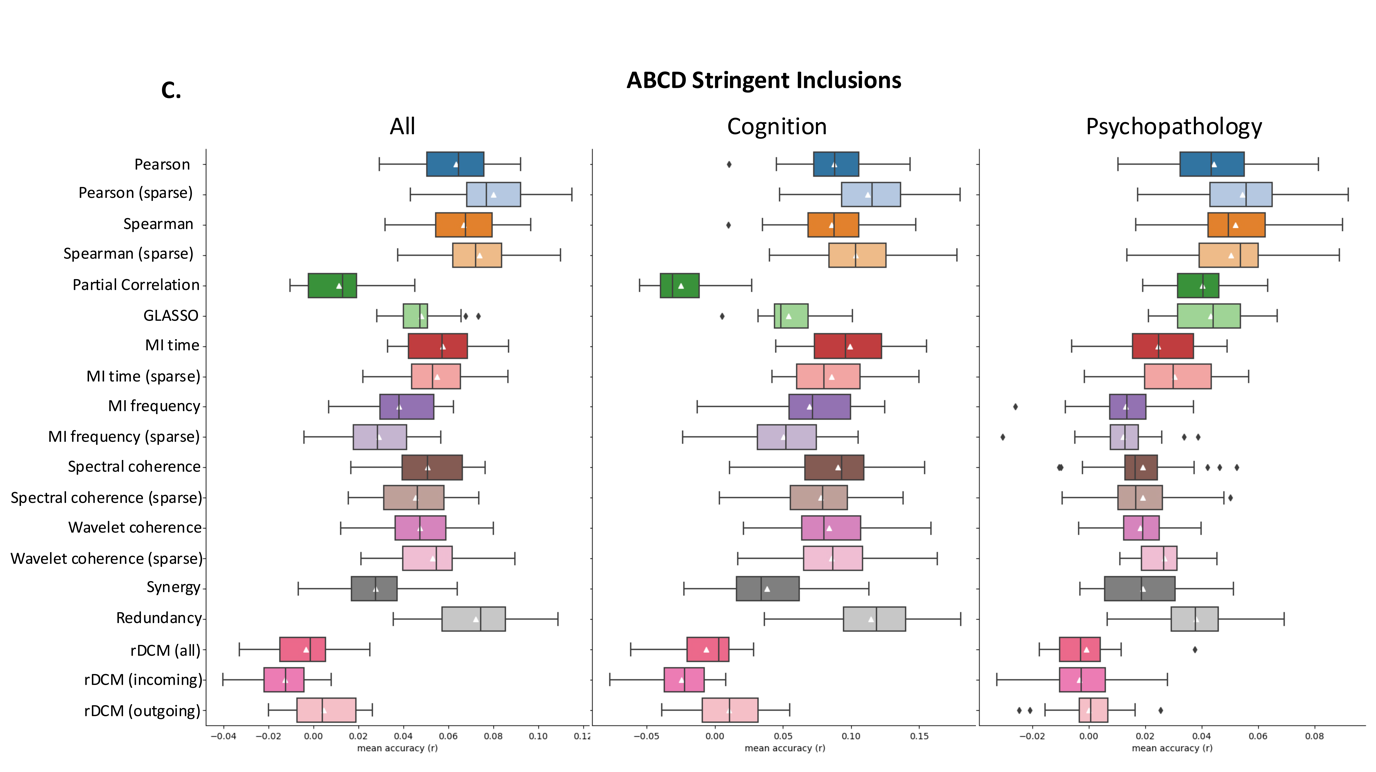
*Supplementary Figure 3.* KRR prediction accuracies presented in the main text, including results after sparsity constraints were imposed. The boxplots show the median and interquartile ranges for accuracies averaged over cross validation folds, repetitions, and behaviours. Mean predictive accuracy for each connectivity measure is shown with a white triangle. Prediction accuracies are shown for HCP participants meeting stringent motion criteria, ABCD participants passing standard quality control inclusion criteria, and ABCD participants meeting stringent motion exclusion thresholds. Measures are ordered as: Correlation → Partial correlation → Mutual Information → Coherence → PID → rDCM.

2. *Behavioural Variable Details*

*Supplementary Table 1.* Behavioural variables used in the KRR for the HCP dataset. For each variable, the corresponding field names in the HCP behavioural csv file are shown together with whether they belonged to the cognition, personality, or other domains. The table has been adapted from Li et al. (2019).

| Description | Name in the HCP dataset | Cognitive | Personality | Others |
| --- | --- | --- | --- | --- |
| Visual Episodic Memory | PicSeq_Unadj_Score | ✓ |  |  |
| Cognitive Flexibility (DCCS) | CardSort_Unadj | ✓ |  |  |
| Inhibition (Flanker Task) | Flanker_Unadj | ✓ |  |  |
| Fluid Intelligence (PMAT) | PMAT24_A_CR | ✓ |  |  |
| Vocabulary (Pronunciation) | ReadEng_Unadj | ✓ |  |  |
| Vocabulary (Picture Matching) | PicVocab_Unadj | ✓ |  |  |
| Processing Speed | ProcSpeed_Unadj | ✓ |  |  |
| Delay Discounting | DDisc_AUC_40K | ✓ |  |  |
| Spatial Orientation | VSPLOT_TC | ✓ |  |  |
| Sustained Attention – Sens. | SCPT_SEN | ✓ |  |  |
| Sustained Attention – Spec. | SCPT_SPEC | ✓ |  |  |
| Verbal Episodic Memory | IWRD_TOT | ✓ |  |  |
| Working Memory (List Sorting) | ListSort_Unadj | ✓ |  |  |
| Cognitive Status (MMSE) | MMSE | ✓ |  |  |
| Sleep Quality (PSQI) | PSQI_Score |  |  | ✓ |
| Walking Endurance | Endurance_Unadj |  |  | ✓ |
| Walking Speed | GaitSpeed_Comp |  |  | ✓ |
| Manual Dexterity | Dexterity_Unadj |  |  | ✓ |
| Grip Strength | Strength_Unadj |  |  | ✓ |
| Odour Identification | Odor_Unadj |  |  | ✓ |
| Pain Interference Survey | PainInterf_Tscore |  |  | ✓ |
| Taste Intensity | Taste_Unadj |  |  | ✓ |
| Contrast Sensitivity | Mars_Final |  |  | ✓ |
| Emotional Face Matching | Emotion_Task_Face_Acc |  |  | ✓ |
| Arithmetic | Language_Task_Math_Avg_Difficulty_Level | ✓ |  |  |
| Story Comprehension | Language_Task_Story_Avg_Difficulty_Level | ✓ |  |  |
| Relational Processing | Relational_Task_Acc |  |  | ✓ |
| Social Cognition – Random | Social_Task_Perc_Random |  |  | ✓ |
| Social Cognition – Interaction | Social_Task_Perc_TOM |  |  | ✓ |
| Working Memory (N-back) | WM_Task_Acc | ✓ |  |  |
| Agreeableness (NEO) | NEOFAC_A |  | ✓ |  |
| Openness (NEO) | NEOFAC_O |  | ✓ |  |
| Conscientiousness (NEO) | NEOFAC_C |  | ✓ |  |
| Neuroticism (NEO) | NEOFAC_N |  | ✓ |  |
| Extraversion (NEO) | NEOFAC_E |  | ✓ |  |
| Emotion Recognition – Total | ER40_CR |  |  | ✓ |
| Emotion Recognition – Angry | ER40ANG |  |  | ✓ |
| Emotion Recognition – Fear | ER40FEAR |  |  | ✓ |
| Emotion Recognition – Happy | ER40HAP |  |  | ✓ |
| Emotion Recognition - Neutral | ER40NOE |  |  | ✓ |
| Emotion Recognition – Sad | ER40SAD |  |  | ✓ |
| Anger – Affect | AngAffect_Unadj |  | ✓ |  |
| Anger – Hostility | AngHostil_Unadj |  | ✓ |  |
| Anger – Aggression | AngAggr_Unadj |  | ✓ |  |
| Fear – Affect | FearAffect_Unadj |  | ✓ |  |
| Fear – Somatic Arousal | FearSomat_Unadj |  | ✓ |  |
| Sadness | Sadness_Unadj |  | ✓ |  |
| Life Satisfaction | LifeSatisf_Unadj |  |  | ✓ |
| Meaning & Purpose | MeanPurp_Unadj |  |  | ✓ |
| Positive Affect | PosAffect_Unadj |  | ✓ |  |
| Friendship | Friendship_Unadj |  |  | ✓ |
| Loneliness | Loneliness_Unadj |  |  | ✓ |
| Perceived Hostility | PercHostil_Unadj |  |  | ✓ |
| Perceived Rejection | PercReject_Unadj |  |  | ✓ |
| Emotional Support | EmotSupp_Unadj |  |  | ✓ |
| Instrument Support | InstruSupp_Unadj |  |  | ✓ |
| Perceived Stress | PercStress_Unadj |  |  | ✓ |
| Self-Efficacy | SelfEff_Unadj |  |  | ✓ |

*Supplementary Table 2.* Behavioural variables used in the KRR for the ABCD dataset. For each variable, the corresponding field names in the ABCD behavioural csv file are shown together with whether they belonged to the cognition or psychopathology domains. The table has been adapted from Li et al. (2019).

| Description | Name in the ABCD dataset | Cognitive | Psychopathology |
| --- | --- | --- | --- |
| Vocabulary | nihtbx_picvocab_uncorrected | ✓ |  |
| Attention | nihtbx_flanker_uncorrected | ✓ |  |
| Working Memory | nihtbx_list_uncorrected | ✓ |  |
| Executive Function | nihtbx_cardsort_uncorrected | ✓ |  |
| Processing Speed | nihtbx_pattern_uncorrected | ✓ |  |
| Episodic Memory | nihtbx_picture_uncorrected | ✓ |  |
| Reading | nihtbx_reading_uncorrected | ✓ |  |
| Fluid Cognition | nihtbx_fluidcomp_uncorrected | ✓ |  |
| Crystalised Cognition | nihtbx_cryst_uncorrected | ✓ |  |
| Overall Cognition | nihtbx_totalcomp_uncorrected | ✓ |  |
| Short Delay Recall | pea_ravlt_sd_trial_vi_tc | ✓ |  |
| Long Delay Recall | pea_ravlt_ld_trial_vii_tc | ✓ |  |
| Fluid Intelligence | Pea_wiscv_trs | ✓ |  |
| Visuospatial Accuracy | lmt_scr_perc_correct | ✓ |  |
| Visuospatial reaction time | lmt_scr_rt_correct | ✓ |  |
| Visuospatial Efficiency | lmt_scr_efficiency | ✓ |  |
| Negative Urgency | upps_y_ss_negative_urgency |  | ✓ |
| Lack of Planning | upps_y_ss_lack_of_planning |  | ✓ |
| Sensation Seeking | upps_y_ss_sensation_seeking |  | ✓ |
| Positive Urgency | upps_y_ss_positive_urgency |  | ✓ |
| Lacks Perseverance | upps_y_ss_lack_of_perserverance |  | ✓ |
| Behavioural Inhibition | bis_y_ss_bis_sum |  | ✓ |
| Reward Responsiveness | bis_y_ss_bar_rr |  | ✓ |
| Drive | bis_y_ss_bas_drive |  | ✓ |
| Fun Seeking | bis_y_ss_bas_fs |  | ✓ |
| Anxious Depressed | cbcl_scr_syn_anxdep_r |  | ✓ |
| Withdrawn Depressed | cbcl_scr_syn_withdep_r |  | ✓ |
| Somatic Complaints | cbcl_scr_syn_somatic_r |  | ✓ |
| Social Problems | cbcl_scr_syn_social_r |  | ✓ |
| Thought Problems | cbcl_scr_syn_thought_r |  | ✓ |
| Attention Problems | cbcl_scr_syn_attention_r |  | ✓ |
| Rule Breaking | cbcl_scr_syn_rulebreak_r |  | ✓ |
| Aggression | cbcl_scr_syn_aggresssive_r |  | ✓ |
| Total Psychosis Symptoms | pps_y_ss_number |  | ✓ |
| Psychosis Severity | pps_y_ss_severity_score |  | ✓ |
| Mania | pgbi_p_ss_score |  | ✓ |
